# The genetic code is not optimized for resource conservation

**DOI:** 10.1101/2021.02.08.430341

**Authors:** Haiqing Xu, Jianzhi Zhang

**Affiliations:** Department of Ecology and Evolutionary Biology, University of Michigan, Ann Arbor, Michigan 48109, USA

## Abstract

Shenhav and Zeevi conclude in a recent article (*Science* 370:683-687) that the universal genetic code (UGC) is optimized for resource conservation because mutations are less likely to increase proteomic nitrogen and carbon uses under the UGC than under random genetic codes (RGCs). Their finding results from miscalculating mutational effects and benchmarking against biased RGCs. Our reanalysis refutes their conclusion.

## Main text

It is well known that environmental nutrients can shape the genomic and proteomic compositions of organisms (*1-5*). Shenhav and Zeevi first corroborated the existence of resource-driven selection in marine microbes (*6*). They then computed the expected random mutation cost (ERMC) in proteomic nitrogen/carbon usage, concluding that the UGC is optimized for nutrient conservation. It is this second part of their study that is conceptually new and the subject of our comment.

In computing the ERMC, Shenhav and Zeevi considered the increase (positive cost) but not decrease (negative cost) of the proteomic nitrogen/carbon content caused by mutations. However, in a nitrogen/carbon-limited environment, if mutations increasing the proteomic nitrogen/carbon content are deleterious, those lowering the content would be beneficial. Therefore, the correct ERMC, referred to as the new EMRC (nEMRC) hereinafter, should be the sum of the positive and negative costs of mutations.

Shenhav and Zeevi reported that the “square” arrangement in the UGC, where nitrogen-rich amino acids are concentrated in one section of the code table instead of being spread over the entire table, reduces its ERMC (*6*). In fact, the “square” arrangement causes mutations to be less likely to increase as well as decrease the proteomic nitrogen usage. Specifically, if a mutation from codon *i* to *j* leads to an increased nitrogen usage, this effect is completely offset by a reverse mutation from *j* to *i*. Let *μ*_ij_, the mutation rate from *i* to *j*, be the probability that a codon *i* is mutated to *j* in a unit time. When *μ*_ij_ = *μ*_ji_ for all codon pairs—assumed in (*6*) and here—and when all codons are equally frequent, the mutational cost measured by nERMC is zero regardless of the structure of the code table or the mutation rate ratio of transitions to transversions (Ts/Tv); that is, the UGC is equally optimized as RGCs in nutrient conservation.

Under unequal codon frequencies, nERMC varies among different code tables, with a mean of zero for RGCs. Regarding the UGC, if the frequencies of codons for nitrogen/carbon-rich amino acids are lower than those for nitrogen/carbon-poor amino acids, for example, as a result of resource-driven selection, mutations will more likely increase than decrease the proteomic nitrogen/carbon content, yielding a positive nERMC or, by Shenhav and Zeevi’s language, a less optimized UGC than RGCs in nutrient conservation.

With the above theoretical consideration in mind, we turned to empirical data. For each of the 39 diverse species examined in (*6*), we computed Pearson’s correlation across codons between the frequency of a codon and the number of nitrogen or carbon atoms in the amino acid encoded by the codon. For nitrogen, the correlation is negative in every species (Fig. 1A), confirming the avoidance of codons encoding nitrogen-rich amino acids (*3*). For carbon, however, the correlation is positive in almost all species (Fig. 1A), suggesting preferential uses of codons encoding carbon-rich amino acids. We thus predict that, compared with RGCs, the UGC will not look optimized in nitrogen conservation but may look optimized for carbon conservation in species with strong codon preferences for carbon-rich amino acids.

**Fig 1.**
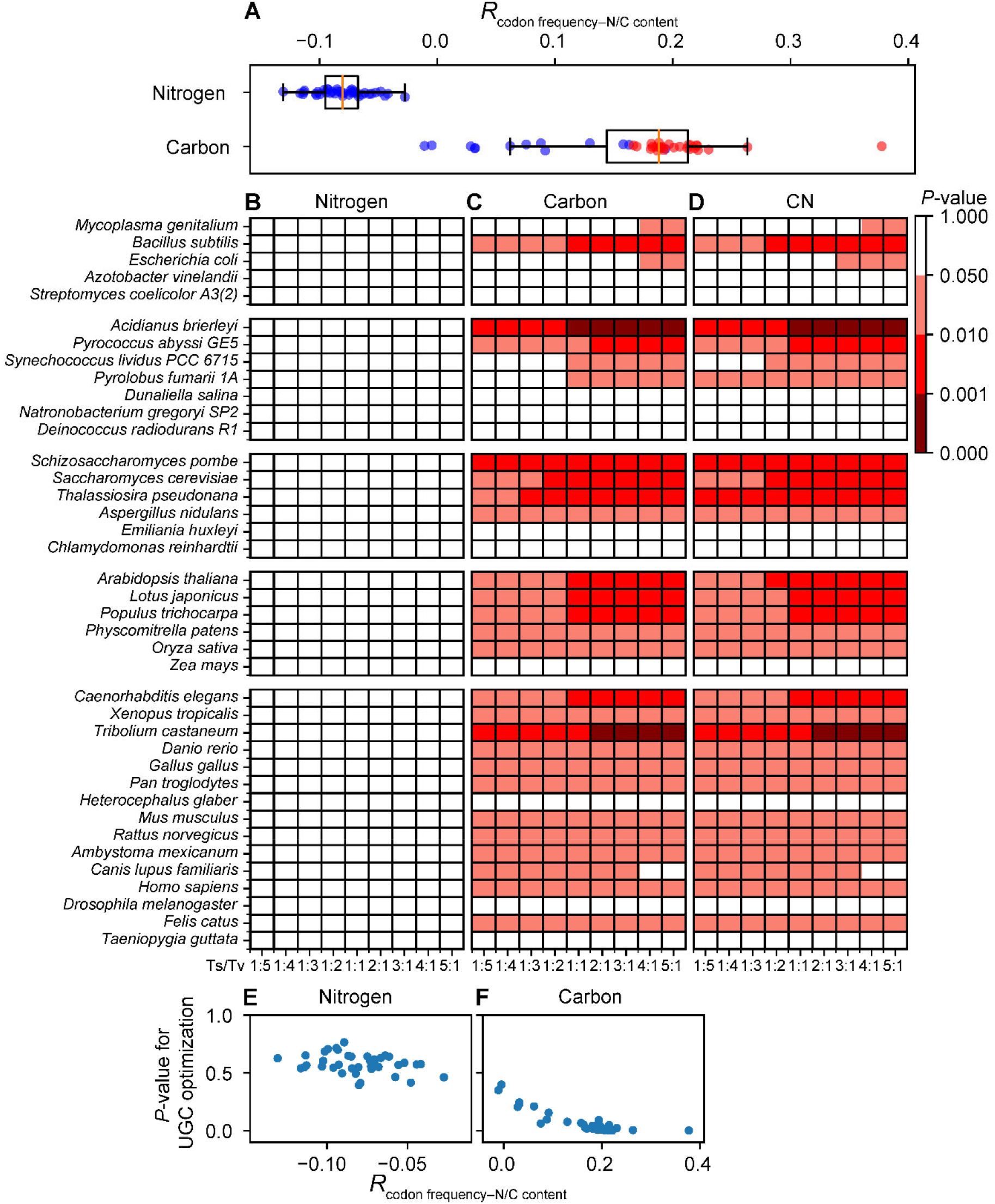
Testing the optimization of the UGC for resource conservation using nERMC. (**A**) Pearson’s correlation (*R*_codon frequency–N/C content_) between the frequency of a codon and the number of nitrogen or carbon atoms of its encoded amino acid in each of 39 species examined. Each dot represents one species. A dot for nitrogen or carbon is marked in red if at least five of the nine examined Ts/Tv values yield significant results in the corresponding species in (B) or (C); otherwise it is marked in blue. The box plot shows the distribution of the 39 data points, with the left and right edges of the box representing the first (qu1) and third (qu3) quartiles, respectively, the vertical line inside the box indicating the median (md), and the whiskers extending to the most extreme values inside inner fences, md ± 1.5(qu3 − qu1). (**B-D**) Heat map of the significance level of the optimization of the UGC for conservation of nitrogen (**B**), carbon (**C**), or both carbon and nitrogen (**D**). Colors indicate nominal *P* values. (**E-F**) Relationship between *R*_codon frequency–N/C content_ and the significance level of the optimization of the UGC for nitrogen (**E**) or carbon (**F**) conservation. The significance level of optimization is determined under Ts/Tv = 3 because Ts/Tv was found to be around 3 in most species (*10*). Pearson’s correlation between *R*_codon frequency–N/C content_ and the significance level of optimization is -0.30 (*P* = 0.063) in (E) and - 0.87 (*P* < 0.001) in (F).

To verify these predictions, we compared the nERMC between the UGC and 1 million RGCs generated following (*6*) under the respective empirical codon frequencies of the 39 species at various Ts/Tv values. Indeed, under no condition does the UGC exhibit a significantly lower nERMC for nitrogen when compared with RGCs (Fig. 1B). Furthermore, the UGC is significantly optimized for carbon (Fig. 1C) and CN (carbon and nitrogen) (Fig. 1D) conservations in some species, although the significance levels are much weaker than originally reported (*6*). As predicted, the UGC is generally less optimized than RGCs for nitrogen conservation (reflected by *P* > 50% in most species in Fig. 1E), and the species exhibiting significant UGC carbon conservations have strong codon preferences for carbon-rich amino acids (Fig. 1A). Clearly, the so-called optimization of the UGC for nutrient conservation (*6*) is unrelated to the origin and evolution of the UGC but a side-effect of codon usage.

Theoretically, even if there were a selection for resource conservation of the UGC that is distinct from the resource-driven selection on codon frequencies, the effects of the two selections would be opposite, because the stronger the latter selection, the less optimized the UGC should be, as reasoned above and verified in the 39 species (Fig. 1E, F). Because the former selection is a second-order selection while the latter a first-order selection and because second-order selections are generally much weaker than first-order selections (*7*), we do not expect the UGC to be optimized for nutrient conservation under these selections. For example, a code-table-altering mutation that reduces the nERMC would lower the nutrient cost of future mutations but increase that of the current proteome to a larger extent, so cannot be fixed.

Another problem is that, when generating RGCs, Shenhav and Zeevi did not allow the number of codons for an amino acid to deviate from that in the UGC, departing from the common practice in testing optimizations of the UGC (*8, 9*). The restriction imposed by the authors limited the diversity of RGCs, potentially biasing the test result. When this restriction in lifted, one can easily design many code tables with lower nERMC than that of the UGC, for example, by assigning only one codon to arginine, the most nitrogen-rich amino acid, instead of six as in the UGC, under the assumption that arginine is indispensable from the proteome. To demonstrate the above point empirically, we applied the commonly used method (*8, 9*) to generate 1 million RGCs and compared them with the UGC in terms of nERMC. In none of the 39 species was the UGC significantly better than the RGCs in nitrogen conservation (Fig. 2A). Similar results were obtained for carbon (Fig. 2B) or CN (Fig. 2C) conservation.

**Fig 2.**
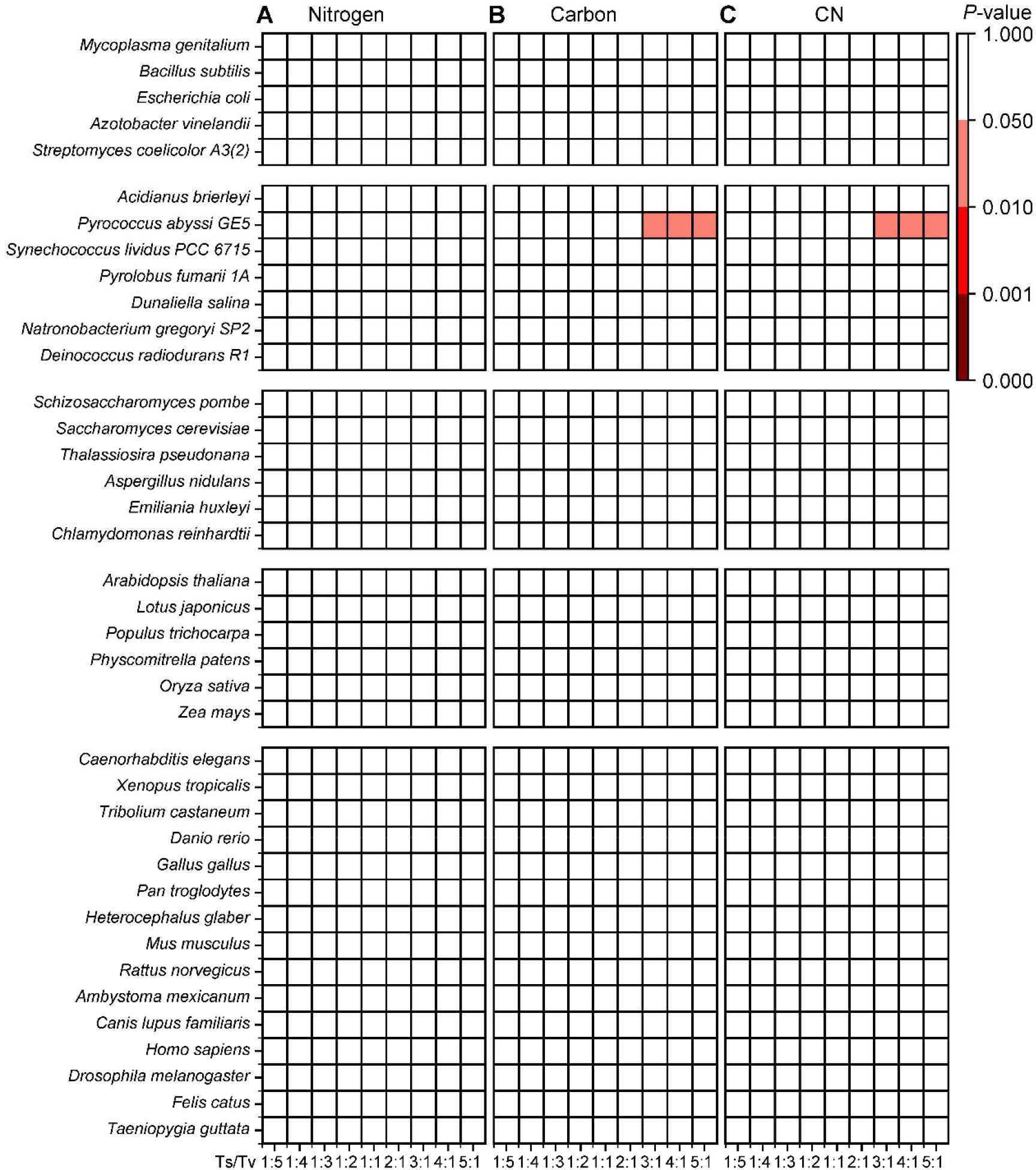
Testing the optimization of the UGC for resource conservation using nERMC and commonly generated RGCs. Heat map of the significance level of the optimization of the UGC for conservation of nitrogen (**A**), carbon (**B**), or both carbon and nitrogen (**C**). Colors indicate nominal *P* values.

In conclusion, Shenhav and Zeevi’s finding of optimization of the UGC for nitrogen/carbon conservation was an artifact of miscalculating mutational effects and benchmarking the UGC against biased RGCs. Their thesis is theoretically untenable and empirically unsupported.

## Acknowledgements

We thank members of the Zhang laboratory for comments. This work was supported by U.S. National Institutes of Health research grant R35GM139484 (J.Z.).

## References

1. J. J. Elser, W. F. Fagan, S. Subramanian, S. Kumar, Signatures of ecological resource availability in the animal and plant proteomes. Mol Biol Evol 23, 1946–1951 (2006).

2. J. J. Elser, C. Acquisti, S. Kumar, Stoichiogenomics: the evolutionary ecology of macromolecular elemental composition. Trends Ecol Evol 26, 38–44 (2011).

3. J. J. Grzymski, A. M. Dussaq, The significance of nitrogen cost minimization in proteomes of marine microorganisms. ISME J 6, 71–80 (2012).

4. D. R. Mende et al., Environmental drivers of a microbial genomic transition zone in the ocean’s interior. Nat Microbiol 2, 1367–1373 (2017).

5. P. M. Berube, A. Rasmussen, R. Braakman, R. Stepanauskas, S. W. Chisholm, Emergence of trait variability through the lens of nitrogen assimilation in Prochlorococcus. eLife 8, e41043 (2019).

6. L. Shenhav, D. Zeevi, Resource conservation manifests in the genetic code. Science 370, 683–687 (2020).

7. D. Graur, A. K. Sater, T. F. Cooper, Molecular and Genome Evolution (Sinauer, Sunderland, Mass., 2016).

8. D. Haig, L. D. Hurst, A quantitative measure of error minimization in the genetic-code. J Mol Evol 33, 412–417 (1991).

9. R. Geyer, A. Madany Mamlouk, On the efficiency of the genetic code after frameshift mutations. PeerJ 6, e4825 (2018).

10. Z. Zou, J. Zhang, Are nonsynonymous tansversions generally more deleterious than nonsynonymous transitions? Mol Biol Evol 38, 181–191 (2021).

